# The Role of Location-Context Binding in Nonspatial Visual Working Memory

**DOI:** 10.1101/352435

**Authors:** Ying Cai, Qing Yu, Andrew D. Sheldon, Bradley R. Postle

## Abstract

Successful retrieval of an item from visual working memory (VWM) often requires an associated representation of the trial-unique context in which that item was presented. We dissociated the effects on fMRI signal of memory load versus context binding by comparing nonspatial VWM for one oriented bar vs. three bars individuated by their location on the screen vs. three items drawn from different categories (orientation, color, and luminance), for which location context was superfluous. Delay-period fMRI signal in frontal and parietal cortex was sensitive to stimulus homogeneity rather than to memory load per se. Behavioral performance revealed a broad range in swap errors, an index of the efficacy of context binding, and subjects were classified as *high swap error* or *low swap error*. During the delay period, the strength of the representation of stimulus location in parietal cortex predicted individual differences in swap errors. During recall, activity in occipital cortex revealed two dissociable neural correlates of context binding: *high swap-error* subjects allocated less spatial attention to the location of the probed item and more spatial attention the location of non-probed items; *high swap-error* subjects also represented the orientation of the probed item more weakly and the orientation of nonprobed items more strongly. Our results suggest context binding is a computation that influences all stages of VWM processing.

**Significance Statement:** Although we often think of the contents of visual working memory (VWM) as representations of the items that need to be remembered, each item’s trial-unique context is also critical for successful performance. For example, if one observes a red, then a black, then a blue car passing through an intersection, vivid memory for the colors, alone, wouldn’t allow one to execute the instruction “Follow the first of the three cars that just drove by.” Although manipulating load is commonly assumed to isolate storage functions, requiring memory for multiple items drawn from the same category also increases demands on the context binding needed to individuate these items. This experiment tracked the influence of context binding on VWM stimulus processing.

## Introduction

Individual differences in human visual working memory (VWM) capacity result from several factors, including the strategic deployment of attention (Fukuda, Woodman, & Vogel, 2015), retrieval-related processes (Unsworth, Fukuda, Awh, & Vogel, 2014), and resistance to interference (Vogel, McCollough, & Machizawa, 2005). In fMRI studies, delay-period activity in the intraparietal sulcus (IPS) scales with the number of items in the memory set before asymptoting at an individual’s VWM capacity (Xu & Chun, 2015; Todd & Marois, 2004, 2005) and has often been assumed to reflect stimulus representation in VWM (Bettencourt & Xu, 2015; Xu, 2017). Here we test the alternative possibility that this activity reflects, at least in part, another computation that is likely to influence VMW capacity. Context binding is the association of each item in the memory set with its unique episodic context. In tests that require VWM for an array of colored squares, for example, the subject must remember not only each of the colors in the array, but also the location at which each color was presented. Intact memory for all the colors but an impaired representation of which had been presented where can lead to “swap errors” (Schneegans & Bays, 2017). Indeed, some theoretical accounts hold that context binding is the factor that determines whether a stimulus can meaningfully be said to be “in” VWM (Oberauer & Lin, 2017).

In one previous study (Gosseries, Yu, et al., 2018), we sought to unconfound the storage demands of a high-load condition from its context-binding demands by varying stimulus category homogeneity within the memory set: Subjects were asked to remember either the direction of motion in one random-dot kinematogram (RDK; *1M* trials), the directions in three RDKs (*3M*), or the direction in one RDK plus the colors of two color patches (*1M2C*). One striking finding was that delay-period activity of the IPS was elevated during *3M* trials relative to both *1M* and *1M2C* trials, suggesting that this activity was sensitive to a factor other than memory load per se. In this experiment, the critical context was ordinal position, because stimuli on three-item trials were presented serially and all in the same location, with the item to be recalled prompted by a digit indicating “1^st^,” “2^nd^,” or “3^rd^.”

In the present experiment, we sought to replicate and extend the results from Gosseries, Yu, et al. (2018) in several important ways. First, we changed the critical contextual dimension from ordinal position to spatial location (by presenting all items simultaneously, in different locations, and indicating the item to be recalled with the location of the memory probe). Would the expected role for IPS in processing location context be similar to or different from its role in processing ordinal context? Furthermore, because the neural correlates of spatial processing are better understood than those of ordinal processing, we could directly assess variation in the representation of stimulus context, as well as in stimulus identity. Second, we sought to remove an important confound in this type of design: A category-homogenous condition like *3M* and a category-heterogeneous condition like *1M2C* do not only differ in terms of demands on context binding, but also differ in other aspects such as level of inter-item interference (Wickens, Born, & Allen, 1963). In this study we took advantage of a broad range of individual differences in swap errors, a hallmark of a context-binding failure that is not expected to be sensitive to interference at the level of stimulus category. Third, instead of using MVPA decoder performance as a proxy for the strength of stimulus representation, we used multivariate inverted encoding modeling (IEM) to obtain interpretable quantitative estimates of the effects of context binding on nonspatial stimulus representation in VWM. Finally, we assessed generality by changing stimulus domains – one orientation vs. three orientations vs. one orientation + one color + one luminance.

## Material and Methods

### Subjects

Eighteen right-handed volunteers (10 females, aged 18-25 years [Mean (SD) = 21.70 (1.75)]) from the University of Wisconsin–Madison community participated in the behavior-only experiment for remuneration ($10/h). Two of the 18 elected not to participate in the fMRI study, and so 16 of these subjects (8 females, aged 18-25 years (Mean (SD) = 20.50 (1.78)) also participated the fMRI experiment ($20/h). The n of 16 was selected to assure sufficient power because a previous study using similar methods found significant effects with n = 12 (Gosseries, Yu, et al. 2018; Bettencourt & Xu, 2015; Xu, 2017). All subjects provided informed consent according to the procedures approved by the Health Sciences Institutional Review Board at the University of Wisconsin–Madison. Subjects had normal or corrected-to-normal vision, no reported history of neurological or psychiatric disease, and, for the fMRI experiment, no contraindications for MRI.

### Stimuli

There were three trial types: delayed recall (a.k.a. “delayed estimation”) of an oriented bar (*1O*), of one from a memory set of 3 oriented bars (*3O*), or of one item from a memory set of 1 oriented bar, 1 color patch, and 1 luminance patch (*1O1C1L*). Oriented-bar stimuli (length, 4°; width, 0.08°) were rendered as the black diameter of a white circular patch. Sample stimuli could appear in one of nine possible orientations ranging from 0–160°, in 20° increments, with a jitter of ±1–5° determined randomly on each trial. Color stimuli were presented on 4°-diameter circles, and drawn from a pool of 9 colors that were equidistant along a circle in CIE L*a*b* color space (L = 70, a = 20, b = 38, radius of 60; sample items were therefore equiluminant, varying mainly in hue and slightly in saturation), with a randomized jitter of ±1–5° on each trial. Luminance stimuli were comprised of a gray annulus (diameter = 2.67°) inside a white ring (RGB values ([0, 0, 0]; diameter = 4°). The annulus could take on one of 9 grayscale values ranging equidistantly from light gray ([0.03, 0.03, 0.03] to darkest gray ([0.97, 0.97, 0.97]), with jitter of ±1–5° determined randomly on each trial.

On all trials, masks were rendered as a white circular patch bisected by 18 black 0.08° x 4° bars, all intersecting at their midpoints and each separated in orientation by 10°.

Recall displays comprised a circular stimulus patch – initially “empty” -- and a response wheel centered on fixation, with a radius to its outer edge of 9.2° and a width of 2°. Varying continuously around the response wheel were all possible values of the category being tested. For orientation, this was rendered as 20 equally spaced black bars (0.05° x 1.8°), ranging in orientation from 0–171°, in 9° increments. For color and luminance, all 180 values of that dimension were evenly distributed along the circle. The angle of rotation of the response wheel varied unpredictably from trial-to-trial, to discourage response planning during the delay. At the onset of the recall display a cursor (a conventional “mouse” arrow) was always positioned at central fixation, and the stimulus patch was rendered with a randomly determined value rendered in the format of the sample stimuli. As soon as the subject began to move the trackball of the response box (see “Behavioral tasks”) the cursor moved correspondingly, and the stimulus patch took on the value corresponding to the location on the response wheel that was nearest to the cursor.

Throughout the experiment, the background screen color was gray [0.5, 0.5, 0.5] (Figure 1).

**Figure 1.**
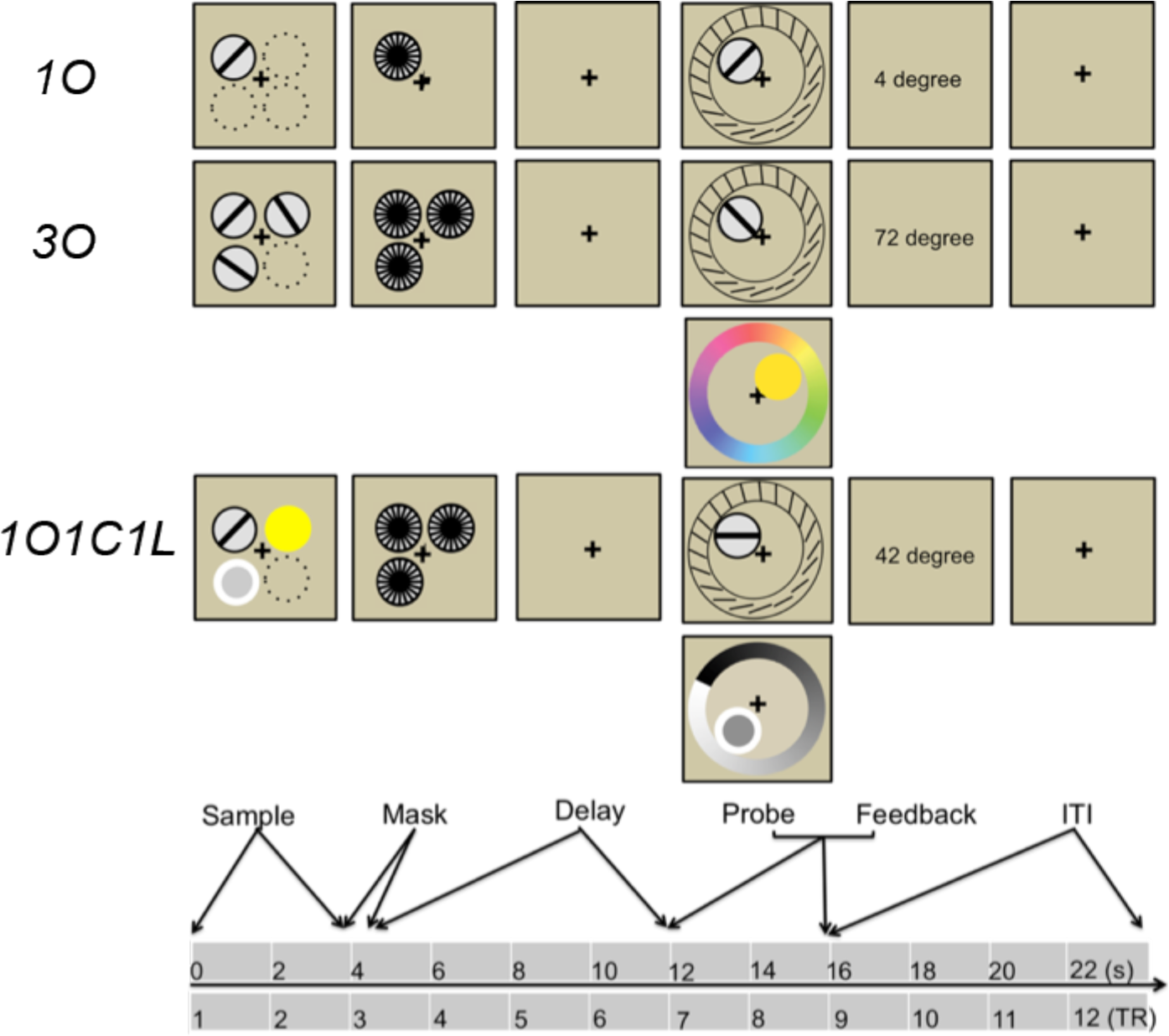
Schematic illustration of the behavioral task. Dotted circles indicate the other possible stimulus presentation locations, but were not presented during experiment.

### Behavioral tasks

Each trial of the *1O* task began with the 4-sec presentation of a sample item equiprobably and unpredictably at one of four possible locations, each in one quadrant of the screen, and each with horizontal and vertical eccentricities from fixation of 5°. The identity of the sample, drawn equiprobably and unpredictably from a pool of 9 orientations, varied independently of location. The 8-s delay period began with a mask presented at the same location as the sample. Responses were made by moving a cursor with a trackball and “clicking” on the recalled orientation with a button press. As soon as the trackball began to move, a bar appeared within the circular patch with an orientation, updating in real-time, that matched the orientation on the wheel that was closest to the cursor. RT was computed as the latency between response-wheel movement onset and button press. Feedback, indicating the error between the recalled orientation and the sample orientation (in degrees) was presented centrally, replacing the fixation cross, appearing immediately after the response until the end of the 4s response window. ITI was 2 s for the behavior-only experiment, and 8 s for the fMRI experiment. A black fixation cross was present at the center of the screen throughout each block of trials, and subjects were instructed to fixate it throughout the block.

*3O* trials followed the same procedure was as *1O*, with the following adjustments necessitated by the greater number of items: on each trial, each of the three sample stimuli was drawn randomly from the pool of 9, without replacement; the three were displayed simultaneously, with each of the four possible configurations of location occurring equiprobably and unpredictably; each of the three stimulus locations was masked; and the location of the probed item occurred equiprobably and unpredictably. *1O1C1L* followed the same procedure was as for *3O*, except that each trial featured one sample item drawn from each of the three stimulus categories, and the category of the probed item occurred equiprobably and unpredictably (Figure 1).

#### Behavior-only experiment

Testing was broken into two blocks of *1O* and *3O* trials and three blocks of *1O1C1L* trials. All blocks contained 50 trials, and block order was counterbalanced across subjects by drawing the first 18 orders from a Latin square. Each block of *1O* and *3O* trials presented 25 of each, in a randomized sequence (thereby yielding a total of 50 *1O* responses and 50 *3O* responses per subject), and each block of *1O1C1L* trials included 17 probes of two of the categories and 16 of the remaining category, randomized within block and balanced across the three blocks, thereby yielding 50 *1O1C1L* orientation responses, 50 *1O1C1L* color responses, and 50 *1O1C1L* luminance responses. All the experimental stimuli were controlled by the Psychophysics Toolbox (http://psychtoolbox.org; Brainard, 1997) running in MATLAB (MathWorks), presented on a 60 Hz projector with a screen width of 32.5cm (iMac). The viewing distance was 62 cm.

#### fMRI experiment

There were two scanning sessions, and during the first session subjects first performed 4 blocks of 3-item trials: 9 trials of *3O* and 9 trials *1O1C1L* in a randomly determined order during each block; 3 probes of each category on *1O1C1L* trials in a randomly determined order during each block. Next, subjects completed 8 18-trial blocks of *1O* trials, with each orientation appearing twice in a randomly determined order during each block. In the second fMRI scanning session, subjects performed an additional 12 18-trial blocks of *1O* trials. (The larger number of *1O* trials was needed to train the IEM models on which data from all trial types would be tested.)

#### Analysis of behavioral data

Reaction time (RT) of the response-ending button press was collected and raw response error distance estimated by the distance on the response wheel between the subjects’ selection and the true target value (in degrees). Trials without responses were excluded. For *1O* and *1O1C1L* trials, response error was fit to a two-factor mixture model that estimated the proportion of responses made to the sample (i.e., the probability of a target response (*p*T), and the probability of guess responses (*p*U), as well as the precision of target responses (κ) (Oberauer & Lin, 2017). For *3O* trials a third factor, the probability of a response to a non-target (*p*N; a.k.a. “swap error”), was included in the model. Parameter estimates were obtained using maximum-likelihood estimation (expectation maximization) using MATLAB routines available at http://www.bayslab.com. Trials lacking a response were excluded from the model fitting process. Differences across trial type in descriptive measures and in model parameters were assessed with repeated measures one-way ANOVAs, and significant effects were followed up with paired *t*-tests. For descriptive measures, we only focused on response error, because the RT was necessarily noisy since it included the time to adjust the response dial with a trackball positioned adjacent to the thigh of the supine subject. For model estimates, we focused on κ and *p*T, because *p*U was highly collinear with *p*T in *1O* and *1O1C1L* trials (i.e., for the two-factor mixture model, *p*U+*p*T=1), and *p*N could only be estimated in *3O* trials.

The *p*N parameter estimated from *3O* trials provides a measure of the efficacy of context binding, because a swap error corresponds to a trial on which the subject has forgotten the location context of the item that s/he is recalling. Post hoc inspection of performance from the behavior-only experiment revealed a clear bimodal distribution in estimates of *p*N, with 6 subjects having a *p*N at or near 0 (indicating effectively no swap errors) and the remaining 10 subjects all having a *p*N of 0.127 or higher (Figure 2A). Based on this pattern in the behavior-only experiment, subjects were grouped into *low swap-error* and *high swap-error* groups for the analyses of the fMRI experiment.

For the behavioral task in the fMRI experiment, RT and response errors were collected in the same way, but model fitting was only conducted for *1O* trials due to the limited number of *3O* and *1O1C1L* trials. Correlations were carried out on all measures of behavioral performance on *1O* trials from the two experiments, with the exception of RT, to assess the stability of subjects’ performance.

### fMRI methods

#### General procedure and behavioral tasks

The fMRI experiment comprised two scanning sessions, each lasting about 1.5 hr. The first of the two scanning sessions followed the behavior-only task by x-y days, and the second scanning session followed the first by 2-28 days. Scanning of 3-item trials preceded scanning of *1O* trials to minimize the likelihood that subjects would process orientation stimuli different from color and luminance stimuli on *1O1C1L* trials. All the experimental stimuli were controlled by the Psychophysics Toolbox (http://psychtoolbox.org; Brainard, 1997) running in MATLAB (MathWorks), presented via a 60 Hz projector (Silent Vision 6011; Avotec) backprojecting onto a screen mounted inside the bore of the scanner, and viewed through a coil-mounted mirror. The viewing distance was 69 cm and screen width was 33 cm.

#### Data acquisition

Whole-brain images were acquired with a 3 Tesla scanner (Discovery MR750; GE Healthcare) at the Lane Neuroimaging Laboratory at the University of Wisconsin–Madison. For all subjects, a high-resolution T1-weighted image was acquired with a fast spoiled gradient-recalled-echo sequence (TR = 8.2 ms, TE =3.2 ms, Flip angle =12°, 160 axial slices, 256 x 256 in-plane, 1 mm isotropic). A T2*-weighted gradient echo pulse sequence was used to acquire data sensitive to the BOLD signal while subjects performed the VSTM task (TR=2000 ms, TE=25 ms, Flip angle = 60°, within a 64 x 64 matrix, 42 sagittal slices, 3 mm isotropic). Each of the twenty fMRI scanning runs generated 213 volumes (excluding disdaqs).

#### Preprocessing

fMRI data were preprocessed using the Analysis of Functional Neuroimages (AFNI) software package (http://afni.nimh.nih.gov; Cox, 1996). All volumes were spatially aligned to the first volume of the first run using rigid-body realignment, then aligned to the T1 volume. Volumes were corrected for slice-time acquisition, and linear, quadratic, and cubic trends were removed from each run to reduce the influence of scanner drift. For univariate analyses, data were spatially smoothed with a 4-mm FWHM Gaussian, and z-scored separately within run for each voxel. For IEM and MVPA analyses (see below), data were z-scored separately within run for each voxel, but were not smoothed. All analyses were carried out in each subject’s native space.

#### Data analysis

*Univariate analyses and ROI creation*. A modified general linear model (GLM) was fit to data from all *3O* and *1O1C1L* trials and from 36 randomly selected *1O* trials. It included regressors modeling the sample presentation, delay, and recall periods with boxcars of 4 s, 8 s, and 4 s, respectively, each convolved with the canonical hemodynamic response function supplied with AFNI.

A different set of anatomically constrained, functionally defined regions of interest (ROI) was created for each of the three categories of analysis: BOLD signal intensity, IEM of stimulus orientation, and MVPA of stimulus location. For all three, anatomical regions were generated from the standard anatomical masks for occipital, parietal, and frontal cortex from the MNI152_T1_1mm template and warping them to each subject’s native space. For BOLD signal intensity analyses, an *Occipital Sample* ROI was generated for each subject by selecting the 400 voxels with the highest *t*-values for the contrast [Sample_*3O*_ – Sample_*1O*_] within anatomically defined occipital cortex, and *Parietal Delay* and *Frontal Delay* ROIs were generated by selecting the 400 voxels with the highest *t*-values for the contrast [Delay_*3O*_ – Delay_*1O*_] within each of these anatomically defined regions. Within each of these ROIs, trial-averaged time series for each of the three trial types were generated and converted to mean percentage signal change from baseline (first TR of the trial), and comparisons of BOLD signals of each trial type versus baseline and differences between trial types were carried out with *t* tests (all *p* values FDR-corrected across TRs, ROIs and comparisons; Figure 3A). For IEM analyses, four sets of “sample location-specific” ROIs were generated with the top 400 voxels within each of the three anatomical regions responding to the contrasts [Sample_upper left_ – baseline], [Sample_upper right_ – baseline], [Sample_lower left_ – baseline], and [Sample_lower right_ – baseline]. Finally, for MVPA analyses, *“location-general”* ROIs were created with the top 400 voxels within each of the three anatomical regions identified with the contrast [(Sample_upper left_ + Sample_upper right_ + Sample_lower left_ + Sample_lower right_) − baseline].

*Task-related patterns of covariation*. We used ANCOVA to evaluate evidence for correlated sensitivity across trial types (specifically, across *1O* vs. *3O* and across *1O1C1L* vs. *3O*) of two of dependent variables -- BOLD signal intensity and behavioral precision – seen to covary in previous studies studies (Emrich, Riggall, Larocque, & Postle, 2013; Grosseries, Yu, et al., 2018). Unlike simple correlations, ANCOVA accommodates the fact that each subject contributes a value for each level of the factor of trial type. It removes between-subject differences and assesses evidence for “within-subject correlation” -- the extent to which variation in one dependent variable can be explained by variation in a second (Bland & Altman, 1995). For analyses including delay-period fMRI activity, the BOLD signal was averaged across TRs 6 and 7, those least likely to be contaminated by Sample-related signal.

*Multivariate inverted encoding modeling*. We estimated population-level neural representations of the orientation of oriented-bar stimuli with multivariate inverted encoding modeling (IEM; Serences & Saproo, 2012; Sprague et al., 2018). To optimize estimation from *1O* trials, four IEMs were trained within each of the three anatomical ROIs, one for each location at which a sample could appear on the screen. The four resultant location-specific reconstructions were averaged prior to assessment of the results of IEM training and testing.

To build our IEMs we assumed that the responses of each voxel can be characterized by activity in 9 hypothesized tuning channels, one corresponding to each of the 9 possible sample orientations. Following previous work(Ester, Sprague, & Serences, 2015; Yu & Shim, 2017), the idealized feature tuning curve of each channel was defined as a half-wave-rectified and squared sinusoid raised to the seventh power. Before feeding the preprocessed data into the IEM, a baseline from each voxel’s response was removed in each run using the following equation from (Brouwer, & Heeger,2011):

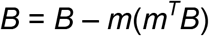

in which *B* represented the data matrix from each run with size *v* × *c* (*v*: the 400 location-specific voxels; *c*: the 9 orientations) and *m* represented the mean response across all stimulus conditions of length *v*. Next, for trials corresponding to each of the four sample locations, we randomly divided the data into a training set (81 trials) and a test set (9 trials; these numbers were selected to match the total number of *1O1C1L* and *3O* trials). We computed the weight matrix (*W*) that projects the hypothesized channel responses (*C_1_*) to actual measured fMRI signals in the training dataset (*B_1_*), and extracted the estimated channel responses (*Ĉ*_2_) for the test dataset (*B_2_*) using this weight matrix. The relationship between the training dataset (*B_1_, v × n, n*: the number of repeated measurements) and the channel responses (*C_1_, k × n*) was characterized by:

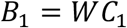

Where *W* was the weight matrix (*v × k*).

Next, the least-squared estimate of the weight matrix 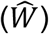 was calculated using linear regression:

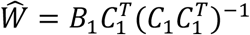

The channel responses (*Ĉ*_2_) for the test dataset (*B_2_*) were then estimated using the weight matrix 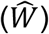:

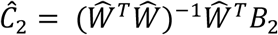

Having thus estimated the weight matrix mapping each voxel’s response to each orientation channel from the training dataset, we inverted this matrix to estimate channel responses on each test trial. The average response output for each channel across trials was obtained by circularly shifting each response to a common center of 0°. To generate smooth, 180-point channel tuning functions (CTF, also referred to as “reconstructions”) we repeated the encoding model analysis 180 times and shifted the centers of the orientation channels by 1° on each iteration (Brouwer, & Heeger, 2009). The CTFs were averaged across permutations and averaged across the four location-specific models. The same weight matrix trained in this fashion on data from *1O* trials was used to reconstruct the neural representation of orientation in *1O, 3O*, and *1O1C1L* trials. More specifically, within each ROI, separate IEMs were trained on *1O* data from each time point in the trial and then tested (i.e., reconstructions attempted) at same time point with data from *1O, 3O*, and *1O1C1L* trials separately. For *3O* trials, the location-specific IEM to be used for IEM testing was assigned according to the orientation to be tested for recall. For *1O1C1L* trials, testing was carried out with the location-specific IEM congruent with the location occupied by oriented-bar in the sample array.

To quantify the results, the CTF in each ROI for each subject was fit with an exponentiated cosine function of the form:

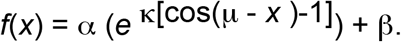

Here, α and β control the vertical scaling (i.e., signal over baseline) and baseline of the function, respectively, and κ and μ control the concentration (the inverse of dispersion) and center of the function, respectively. No biases in reconstruction centers were expected or observed, so we fixed μ at 0. Fitting was performed by combining a GLM with a grid search procedure. We first defined a range of plausible κ values (from 1 to 30 in 0.1 increments). For each possible value of κ, we generated a response function using the fitting equation after setting α to 1 and β to 0. Next, we generated a design matrix containing the predicted response function and a constant term (i.e., a vector of *1s*) and used ordinary least-squares regression to obtain estimates of α and β (defined by the regression coefficient for the response function and constant term, respectively). We then selected the combination of κ, α and β that minimized the sum of squared errors between the observed and predicted reconstructions.

We assessed the significance of CTF amplitude and precision using a bootstrapping procedure in which, for each ROI, we randomly selected, with replacement, 16 CTFs from the pool of 16 subjects being tested, and averaged them. This step was repeated 2500 times, yielding 2500 unique stimulus reconstructions. We then estimated the amplitude of each reconstruction and a *p* value was computed as the proportion of permutations for which amplitude estimates ≤ 0 was obtained (Ester et al., 2015; Ester, Sutterer, Serences, & Awh, 2016). For these and all subsequent bootstrapping analyses, all the *p* values were FDR-corrected across ROIs, TRs, and tasks.

Next, we assessed of the influence of individual differences in context binding on the neural representation of orientation by assessing orientation CTFs for the *low swap-error* group and the *high swap-error* group. These analyses focused on the TRs and ROIs for which CTFs in multi-item trials were reliable at the (n=16) group level, and they were carried out for both the amplitude and precision parameters. First, to assess CTFs versus baseline, we carried out the bootstrapping procedures separately for each group. Next, to compare between groups, the CTF estimate from each permutation for *high swap-error* group was subtracted from the corresponding value for the *low swap-error* group. One-tailed tests were conducted with *p* referring to the proportion of the 2500 subtractions with a value ≤ 0.

Finally, one would expect that individual differences in *p*N, a behavioral measure, would be reflected in neural evidence for inappropriate activation of non-probed items. To assess this prediction, we operationalized “orientation recall specificity” as the difference between probe-epoch CTFs of the probed versus of a non-probed item, and compared this measure between swap-error groups. To compute *orientation recall specificity* for *3O* trials, for each trial one of the two non-probed sample orientations was selected at random; To compute this measure for *1O1C1L* trials, the measure of “probed” CTFs was taken from the 1/3 of trials on which the orientation stimulus was probed, and the measure of “non-probed” CTFs (of orientation) was taken from 1/2 of the trials (randomly selected) on which color was probed and from 1/2 of the trials (randomly selected) on which luminance was probed. For these analyses, all reconstructions were carried out with location-congruent IEMs, and significance of all comparisons was assessed with bootstrapping.

*Decoding the representation of stimulus location*. To examine the neural representation of sample location (which can also be construed as covert spatial attention), we trained MVPA classifiers on data from 18 randomly selected *1O*-trial runs to discriminate among the four sample locations, and tested performance on the held-out *1O* trials (to match the number of *3O* and *1O1C1L* trials), the *3O* trials, and the *1O1C1L* trials, in each of the *location-general* ROIs. Classification was performed using L2-regularized logistic regression with a lambda penalty term of 25, implemented with the Princeton Multi-Voxel Pattern Analysis toolbox (www.pni.princeton.edu/mvpa/) and custom routines in MATLAB. Training and testing were carried out for each TR (i.e., along the diagonal of a training-testing matrix), the classifier producing a probability estimate (from 0.0 −1.0) of the extent to which the observed pattern on the tested trial matched the trained pattern for each of the four locations. Significance of classifier performance was determined using one-tailed, one-sample *t* tests, testing against chance performance of 0.25.

For *3O* and *1O1C1L* trials, decoding was carried out in two ways. First, we labeled trials according to the location that would be probed at the end of the trial. The results of this analysis were likely to be noisy due to random trial-by-trial variation in which two of the remaining three locations were occupied. Therefore, in a second analysis we labeled trials according to the unique location that was unoccupied (i.e., “empty”) in the sample array, and tested whether the decoding accuracy for the empty location was significantly lower (as assessed by one-tailed, one-sample *t* test) than chance. The second analysis was expected to be more sensitive to detect the representations of location information when multiple locations were occupied.

Finally, to explore possible relations between individual differences in the allocation of spatial attention and in swap errors, we operationalized “location recall specificity” -- a measure analogous to *orientation recall specificity* -- by calculating the difference between probe-epoch MVPA decoding of probed versus non-probed locations. For each *3O* and *1O1C1L* trial, the non-probed location was selected randomly. The bootstrapping statistical methods described above were used to estimate the significance of the differences between the two swap-error groups, and any significant effects were followed-up with group-level correlations between *location recall specificity* and *p*N.

## Results

### Behavior

#### Behavior-only experiment

Here, we report results from only the 16 subjects who also participated in the fMRI experiment. An average of 2.286 trials (SD = 2.920); 4.857 trials (SD = 2.824), and 1.857 trials (SD = 2.824) per subject were excluded in *1O, 3O*, and *1O1C1L* tasks, respectively. Descriptive statistics suggested that task difficulty increased from *1O* to *1O1C1L* to *3O*, as reflected in the mean response error (*F*(2,30) = 41.830, *p* < 0.0001; paired t-test, ts > 3.960, *ps* < 0.002; Table 1). Results from mixture modeling mirrored this pattern, with *p*T highest for *1O*, followed by *1O1C1L* and *3O* (F(2,30) = 16.791, *p* < 0.0001), paired t-test ts > 2.644, *ps* < 0.018; Table 1). Recall precision (κ) also differed across trials types (*F*(2,30) = 16.458, *p* < 0.001), being significantly different between *3O* and *1O* (*t*(15) =5.253, *p* <0.0001) and between *3O* and *1O1C1L* (*t*(15) = 4.325, *p* < 0.001) *trials*, although not differing between *1O1C1L* and *1O* trials (*t*(15) = 1.387, *p* = 0.186). Finally, although the group mean *p*N (swap errors) on *3O* trials was 0.120 (SD = 0.116), 6 subjects had a *p*N at or near 0 (all *p*Ns < 0.006), indicating that these subjects made effectively no swap errors, whereas the remaining 10 subjects all had a *p*N of 0.127 or higher, corresponding to an average of nearly 20% swap errors on *3O* trials for these ten subjects (Figure 2A). Based on this pattern, subjects were grouped into *low swap-error* and *high swap-error* groups for fMRI analyses. After subjects were classified as *low swap-error* or *high swap-error* we assessed whether the two groups differed according to other behavioral parameters with two-way mixed ANOVAs. For κ, there were no differences (*F*s(1,14) < 0.731, n.s.). For *p*T, a significant interaction of group by trial type (F(1,14) = 17.753, *p* = 0.001) was followed-up with post-hoc two-sample *t*-tests that confirmed that *p*T was only lower for *3O* in the *high swap-error* group (*t*(14) = 4.085, *p* = 0.001; Figure 2B), a result that follows from the difference in *p*N used to define the two groups.

**Table 1.**
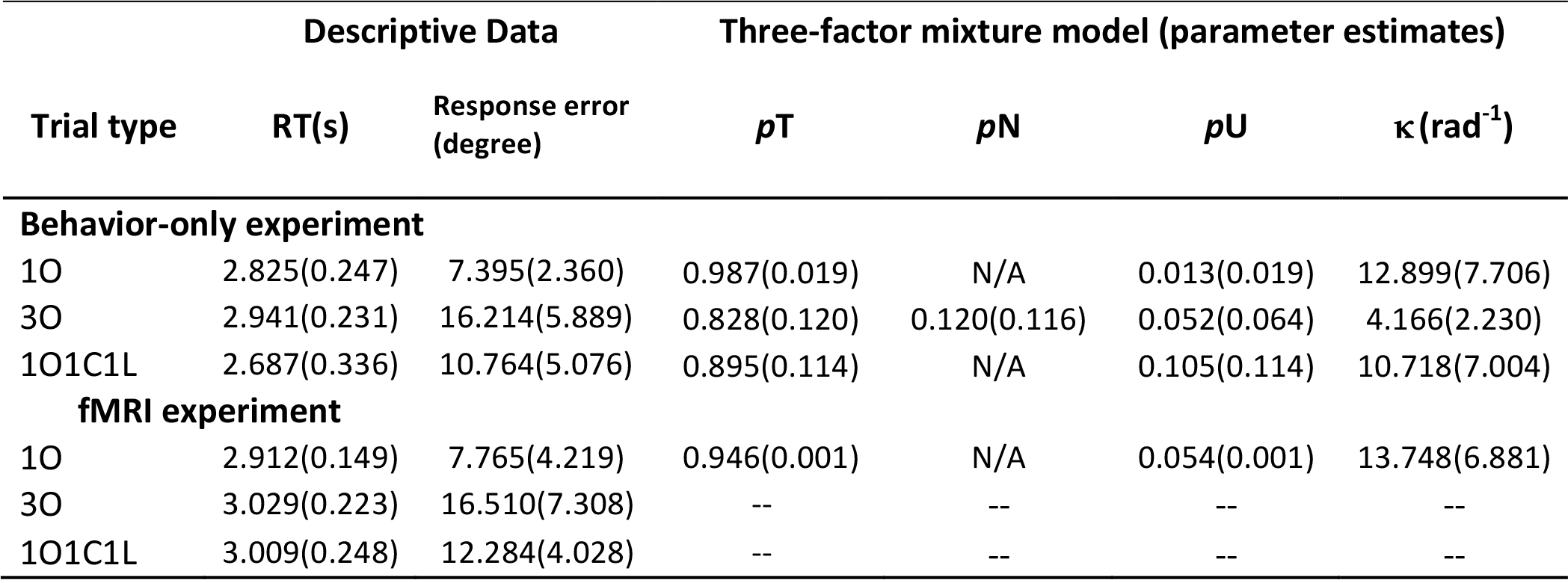
Behavioral results.

#### fMRI experiment

*Behavior during scanning*. An average of 4.62 (SD = 2.07) *1O* trials was excluded, no trials for any subject for either of the other two trial types were excluded. Behavioral performance on *1O* trials performed during scanning was highly correlated with pre-scan testing across the 16 subjects who participated in both (*rs* > 0.841, *ps* < 0.001; Table 1).

**Figure 2.**
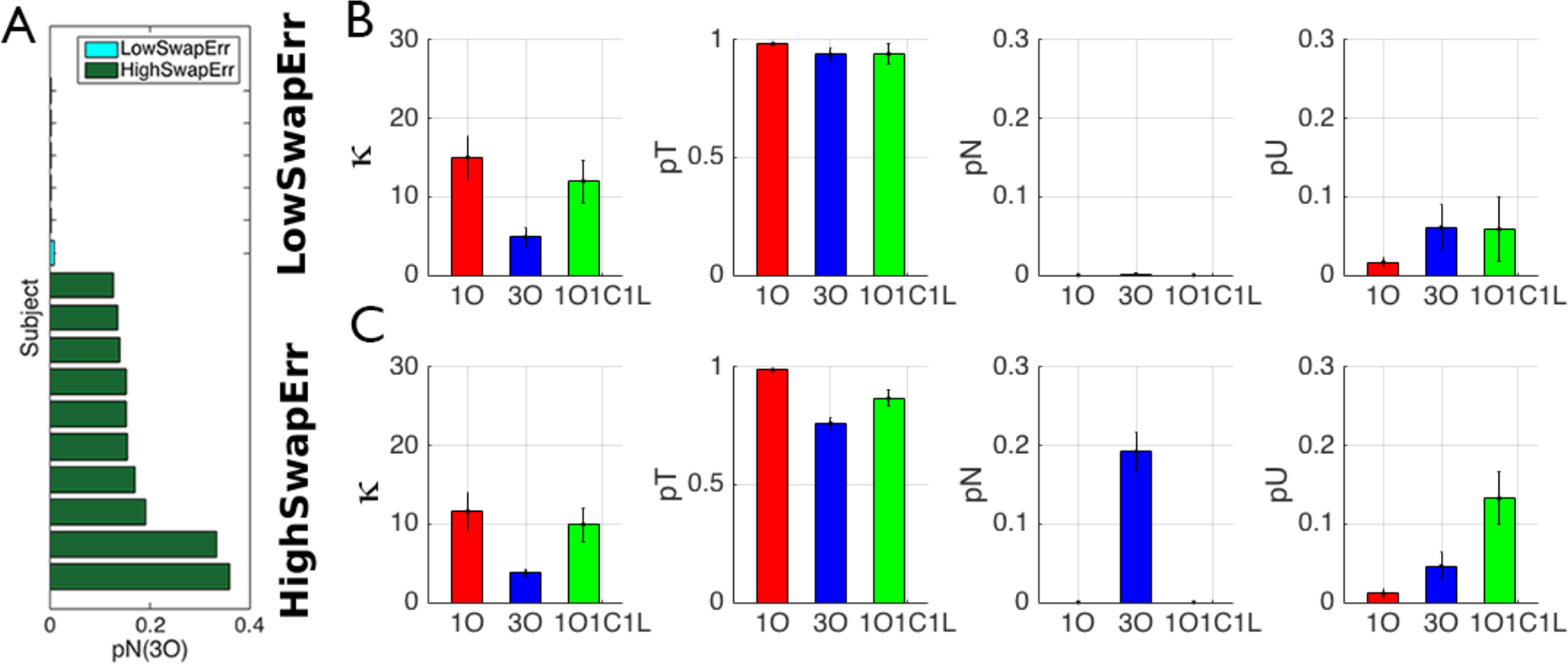
Results from behavior-only experiment. A. Distribution of swap errors (pN) across the 16 subjects participating in the fMRI experiment, sorted by magnitude, and color coded to indicate classification into the *low-swap error* or *high-swap error* group. B-C. Parameters from mixture modeling, by swap-error group.

### fMRI results

#### BOLD signal intensity

*Occipital cortex*. Trial-averaged signal on all three trial types showed the expected sample-related increase, return to baseline by the end of the delay period, and probe/recall-related increase (IEM; Serences & Saproo, 2012; Sprague et al., 2018). BOLD signal intensity did not differ between *3O* and *1O1C1L* trials during TRs corresponding to the encoding and delay epochs (from TR2-TR8, *ts* < 1.604, *ps* > 0.130), but was higher on *3O* than *1O1C1L* trials during the peak response to probe/recall (TR9-11, *ts* > 2.524, *ps* < 0.048). BOLD signal from all three trial types did not differ from each other or from baseline at TR7 corresponding to late delay (*ts* < 0.806, *ps* > 0.433; Figure 3A). These same patterns were observed when the data were broken out into the two swap-error groups.

The ANCOVA relating delay-period BOLD signal intensity to behavioral precision revealed no significant within-subject correlations between either *1O* and *3O* trials or *1O* and *1O1C1L* trials (Figure 3B).

*Parietal and frontal cortex*. In both regions, trial-averaged signal on all three trial types remained elevated across the duration of the trial, with sample- and delay-related activity greater for *3O* trials than for the other two trial types (from TR3 to TR8, *t*s > 3.041, *p*s < 0.019), and sample-related activity for *1O1C1L* trials greater than for *1O* trials during the encoding epoch (TRs 3-4, *ts* > 2.514, *ps* < 0.048). Beginning with TR6 in the delay period, however, BOLD signal no longer differed between *1O1C1L* and *1O* (*ts* < 1.848, *ps* > 0.138). That is, delay-period activity in parietal and frontal cortex was not sensitive to memory load (one item vs. three items) per se, but was sensitive to stimulus category homogeneity (operationalized as *1O1C1L* vs. *3O*; Figure 3A). These same patterns were observed when the data were broken out into the two swap-error groups. Follow-up analyses indicated that differences in delay-period signal (at TRs 6-7) between neither *1O* and *3O* nor *1O1C1L* and *3O* trials correlated with swap errors across subjects (*ps* > 0.524).

ANCOVAs relating delay-period BOLD signal intensity to behavioral precision revealed significant within-subject correlations between *1O* and *3O* trials in both regions, (*rs* > 0.539; *ps* < .05), and a nonsignificant trend in this direction between *1O1C1L* and *3O* trials in parietal cortex (*r* = 0.458; *p* = .06. Figure 3B).

**Figure 3.**
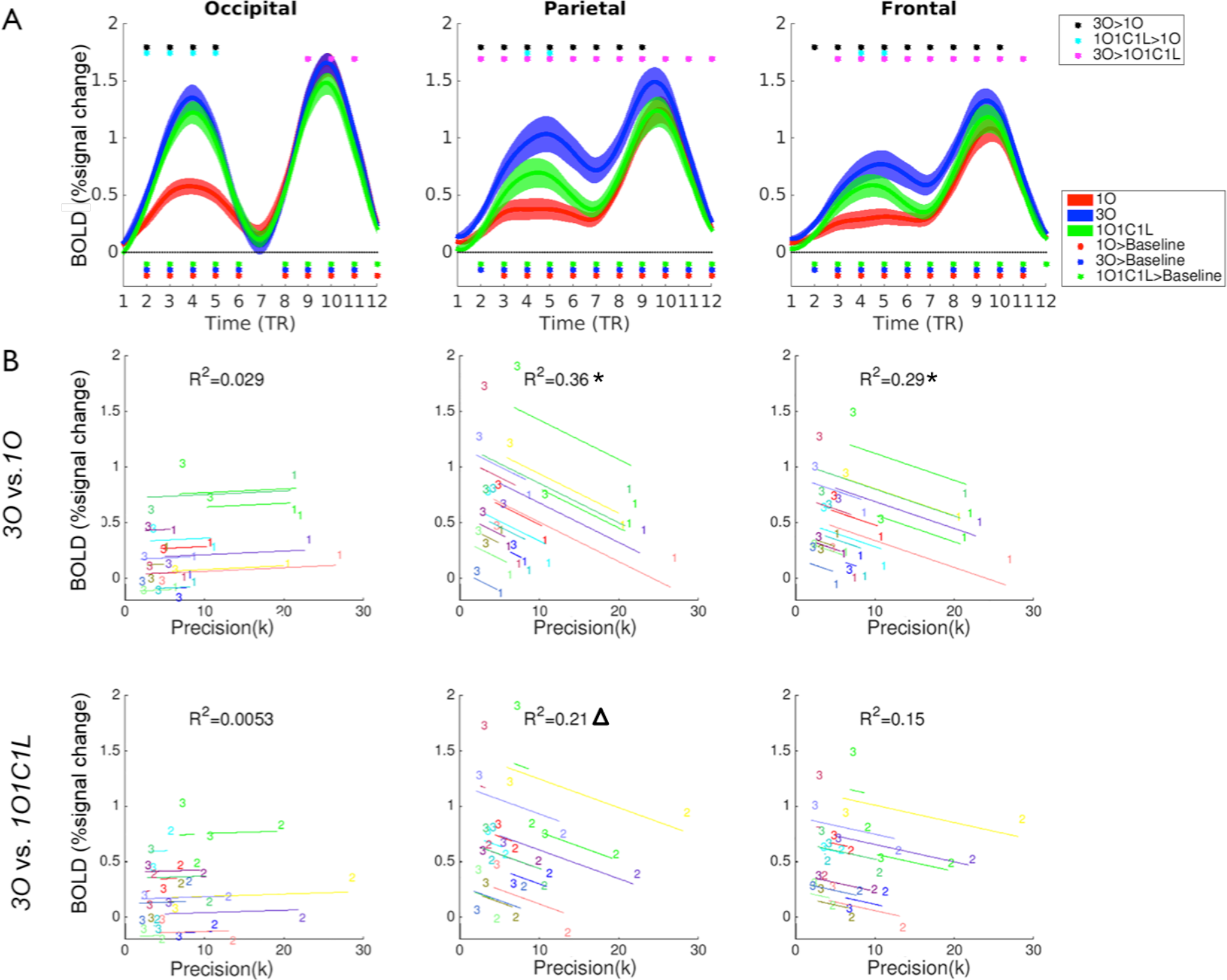
BOLD signal intensity results from three ROIs. A. Trial-averaged BOLD signal. Dots below the x-axis indicate significance vs. baseline; dots above the plots indicate the significance between trial types. B. Within-subject correlations (ANCOVA) between delay-period BOLD signal intensity and behavioral precision of recall. In each plot, data from each subject are portrayed in a different color. The “1,” “3” and “2” symbols indicate individual values in the 1O, 3O, and 1O1C1L tasks, respectively. Asterisks indicate significant correlation at p < 0.05, triangles indicate trends (.05 < p <0.1).

#### IEM of orientation

*Occipital cortex (n = 16)*. On *1O* trials, the amplitude of orientation CTFs was significant during TRs 3-5, spanning sample presentation through early delay (*ps* <0.006), and again during TR10 of the recall epoch (*p*= 0.050; Figure 4A). On *1O1C1L* trials, reconstructions of orientation on these TRs were significant (*ps* < 0.028), and were numerically, but not statistically, smaller than those from *1O* trials (*ps* > 0.252). (Note that for *1O1C1L* trials, testing with data from TR 8 and later only included the 1/3 trials on which orientation was probed). On *3O* trials, although no reconstructions at these TRs were significantly different from baseline, those from TRs 3-5 did not differ from *1O1C1L* trials. At TR10, the CTF amplitude was greater on *1O1C1L* trials than on *3O* trials (*p* = 0.048). No significant effects were found for the concentration parameter for any comparison of IEM reconstructions, here or for any subsequent analyses.

*Parietal and frontal cortex* (*n = 16*). In parietal cortex, IEM reconstructions of orientation were only reliable on *1O* trials, and only during TRs 4-5 (*ps* < 0.005; *ps* > 0.304 for all TRs for *1O1C1L* and *3O* trials). In frontal cortex, the IEM reconstructions of orientation were not significant at any TR for any trial type (*ps* > 0.146). (Note that when IEM of *1O* trials was carried out with a different procedure – leave-one-trial-out cross-validation with all 360 trials – orientation reconstruction was robust across all three trial epochs in occipital, parietal, and frontal cortex. However, because these reconstructions cannot be meaningfully compared with *1O1C1L* and *3O* reconstructions, due to the reduced number of trials in the latter, no results from this more sensitive method for reconstructing orientation from *1O* trials are reported here.)

*Comparison of swap-error groups in occipital cortex*. We assessed the influence of individual differences in context-binding on the neural representation of stimulus orientation in the ROI (occipital) and at the TRs (3-5, 10) for which reconstructions at the group level were most robust (Figure 4A). Beginning with the encoding/early delay TRs 3-5, for *1O* trials, reconstructions were reliable for both groups (*ps* < 0.048), and did not differ between groups (*ps* > 0.094; Figure 4B). For *3O* trials, for the *low swap-error* group, the CTF was significant at TR 3 (*p* = 0.013), trending for TR 4 (*p* = 0.072) and not significant at TR 5 (p = 0.479), and although no CTFs were significant for the *high swap-error* group (*ps* >0.381), the two groups did not differ significantly at any TR (*ps* > 0.241). For *1O1C1L* trials, CTFs for the *low swap-error* group were significant at each of the three TRs (*ps* <0.013), and different from the *high-swap error* group (for which no CTFs were significant; *ps* > 0.070) at TR 5 (*p* = 0.028; Figure 4B).

At TR 10, although the *1O* CTF was only significant for the *low swap-error* group (p < 0.001), the two groups did not differ statistically (*p* = 0.220). For *3O* trials, for the *low swap-error* group there were trends toward a positive reconstruction for the probed orientation and toward a negative reconstruction for non-probed orientations (*ps* < 0.089), a pattern that yielded a significant *orientation recall specificity* effect (*p* = 0.038). For the *high swap-error* group, none of these effects approached significance. The difference in *orientation recall specificity* between the *low* vs. *high swap-error* groups was significant (*p* = *0.031*; Figure 4C).

For *1O1C1L* trials, for the *low swap-error* group, a robust CTF for the probed orientation (*p* < 0.001) and a trend toward a negative reconstruction for the non-probed item (*p* = 0.078) produced a strong *orientation recall specificity* effect (*p* < 0.001). For the *high swap-error* group, the reconstruction for both probed and non-probed items was significant (*ps* < 0.042), and their difference also yielded a significant *orientation recall specificity* effect (*p* = 0.028). *Orientation recall specificity* between the *low* vs. *high swap-error* groups for *1O1C1L* trials was significant (*p* = 0.003; Figure 4D).

**Figure 4.**
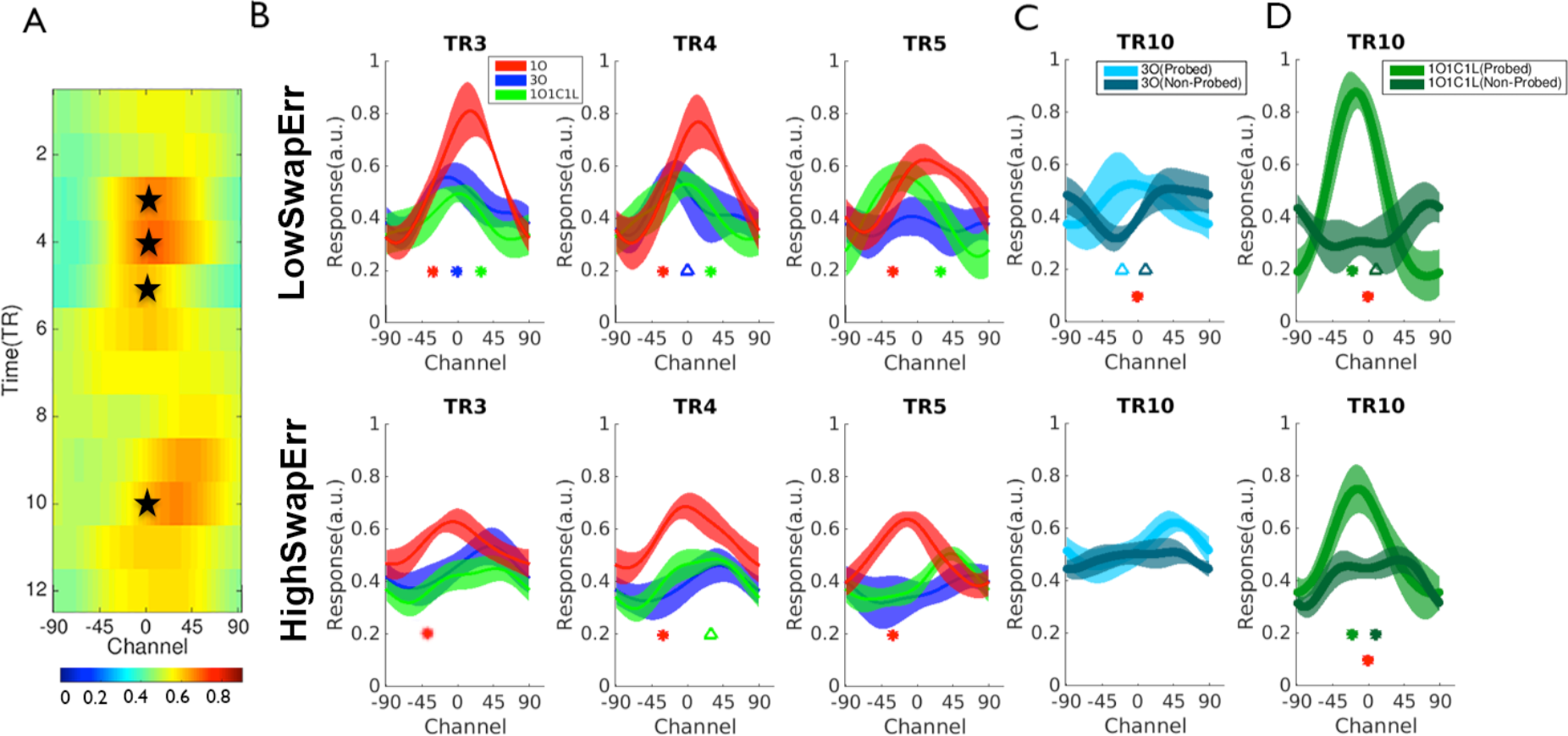
Neural reconstructions of orientation in occipital cortex. A. Group-level (n = 16) IEM reconstructions from *1O* trials, at each TR. Black stars indicate TRs with significant reconstructions. Color scale indicates amplitude of IEM reconstructions (a.u.). B. Reconstructions during encoding/early delay, for the three trial types, for the *low swap-error* group (top row) and *high swap-error* group (bottom row). Color-coded asterisks indicate significant reconstruction at *p* < 0.05, and triangles indicate trends (.05 < *p* < 0.1). C-D. Reconstructions for probed and non-probed items during the probe epoch for *3O* (C) and *1O1C1L* (D) trials. The blue and green asterisks indicate significant neural reconstructions at *p* < 0.05, and triangles indicate trends (.05 < *p* < 0.1). Red asterisks indicate significant *orientation recall specificity* effects at *p* < 0.05.

To summarize this section, during encoding/early-delay portions of the trial, although the neural representation of the to-be-remembered stimulus dimension was quantitatively weaker for the *high swap-error* group, these differences only achieved significance for one condition at one TR (*1O1C1L* trials at TR 5). During recall (TR 10), reconstruction of the probed item was quantitatively stronger for the *low swap-error* group on all trial types, and *orientation recall specificity* was significantly higher for the *low swap-error* groups for both 3-item trial types.

#### Decoding the representation of stimulus location

*Superior location decoding in occipital than in parietal or frontal cortex*. In occipital cortex, MVPA decoding of the probed location was significant for all three trial types across the duration of the trial (TRs 2-12, *ps* < 0.004; Figure 5A). (Note that for *3O* and for *1O1C1L* trials, decoding was most robust during the probe epoch, during which time a stimulus patch occupied the probed location). In parietal cortex, location decoding failed at all TRs using conventional scoring of classifier performance. Re-scoring classifier performance in relation to the empty location in the sample array, however, revealed statistically reliable location information on TRs 3-5 during *3O* trials and on TRs 4-8 during *1O1C1L* trials (*ps* < 0.045; Figure 5A). In frontal cortex, location decoding was above chance at only one TR for only one trial type when MVPA performance was scored according to the empty location (TR 6 on *1O1C1L* trials; *p* = 0.014; Figure 5A), and so no additional analyses were carried out in frontal cortex.

*Differences in MVPA decoding between swap-error groups*. In occipital cortex, decoding accuracy for the probed location did not differ between groups for any trial type for the TRs encompassing the sample and delay epochs (TRs 1-8; *ps* > 0.278). For *1O* trials, location MVPA was superior for the *low swap-error* group for TRs 9-10 of the probe/recall epoch (*p*s < 0.017).

For *3O* trials, for TRs 9-10, decoding of the probed location was significantly higher for the *low swap-error* group (*ps* < 0.010), and decoding of the non-probed location was significantly lower for the *low swap-error* group (*ps* < 0.024), a pattern yielding a significant difference in *location recall specificity* between the two groups (*p* < 0.006, Figure 5B). This finding was followed up with a Spearman correlation indicating that individual differences in *location recall specificity* predicted individual differences in *p*N (*r*(15) = −0.526, *p* = 0.038, Figure 5D).

For *1O1C1L* trials the pattern was qualitatively similar to that from *3O* trials, with the groups differing significantly for decoding of the probed location at TR 10 (*p* = 0.027), and *location recall specificity* significantly higher for the low swap-error group (*p* = 0.018, Figure 5B). For *1O1C1L* trials, Spearman correlation also indicated that individual differences in *location recall specificity* predicted individual differences in *p*N (*r*(15) = −0.506, *p* = 0.048, Figure 5E).

In parietal cortex, classifier performance in relation to the empty location in the sample array was significantly lower for the *low swap-error* group at TRs 4, 6, and 7 on *3O* trials (*ps* < 0.039), and at TRs 5-6 on *1O1C1L* trials (*ps* < 0.031, Figure 5C). For *3O* trials, Spearman correlation relating individual differences in decoding performance against *p*N was nonsignificant for TR 4 (*r*(15) = 0.416, *p* = 0.109; not shown) but significant for TRs 6-7 (*r*(15) = 0.531, *p* = 0.034; Figure 5F). For *1O1C1L* trials, the same correlation collapsed across TRs 5-6 was not significant (*r*(15) = 0.377, *p* = 0.142; not shown).

To summarize, in occipital cortex location MVPA did not differ between swap-error groups across encoding and delay portions of the trial, but was superior for the low-swap error group for all three trial types during recall, and, across all subjects, on both 3-item trial types, individual differences in *location recall specificity* were significantly related to individual differences in *p*N. In parietal cortex, in contrast, location decoding was superior for the *low swap-error* group during late-encoding/early-delay portions of the trial, with a significant group-level correlation of location MVPA and *p*N at TRs 6-7 on *3O* trials.

*Relating spatial processing across networks and across time*. Analyses relating location processing to swap errors on *3O* trials emphasized delay-period effects in parietal cortex and probe-related effects in occipital cortex. To assess evidence for a link between these two sets of observations, we carried out a Pearson correlation of parietal MVPA at TR 6-7 against *location recall specificity* in occipital cortex at TR 10, and found significant evidence that the two were related (*r*(15) = 0.551, *p*= 0.027. Figure 5G). A second analysis with data from *3O* trials from occipital cortex at TR 10, however, failed to find a significant Pearson correlation between *location recall specificity* (as illustrated in Figure 5B) and orientation recall specificity (as illustrated in Figure 4C. *r*(15) = 0.351, *p* = 0.183)).

**Figure 5.**
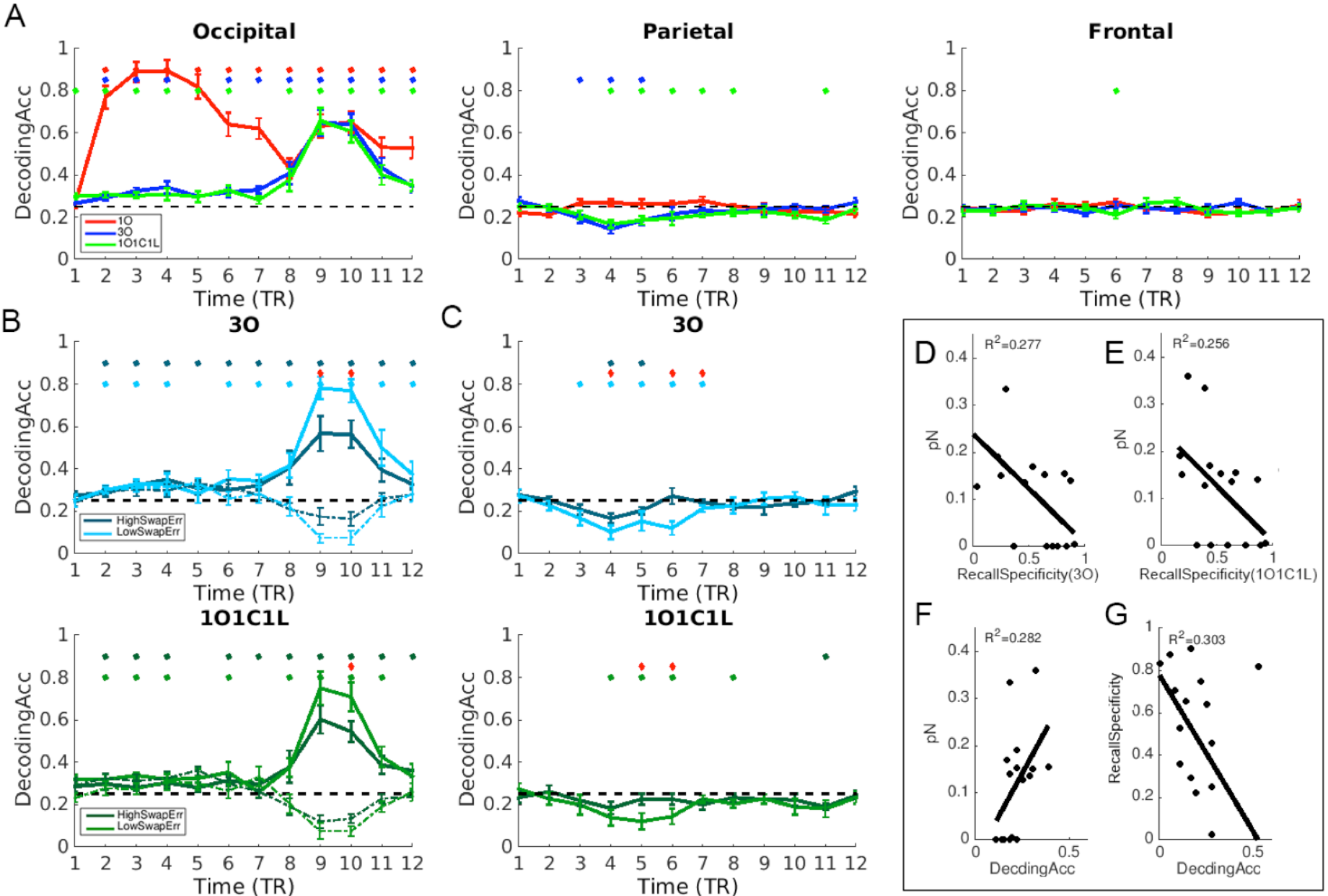
MVPA of location. A. In occipital cortex classifier performance is displayed as decoding accuracy in relation to the probed location during the probe/recall period, with higher values implying stronger representations; in parietal and frontal cortex classifier performance is displayed as decoding accuracy in relation to the empty location in the sample array, and so lower values imply stronger representations. The dashed line indicates chance-level decoding (25%). Color-coded asterisks indicate significant decoding at p < 0.05. B. MVPA decoding in occipital cortex for the probed location (solid line) and for a nonprobed location (dashed line), by swap-error group, in *3O* (top row) and *1O1C1L* (bottom row) trials. C. MVPA decoding in parietal cortex in relation to the empty location on *3O* (top row) and *1O1C1L* (bottom row) trials in the two swap-error groups. For B. and C., color-coded dots indicate significant decoding of the probed location at *p* < 0.05; red asterisks indicate significant differences in *location recall specificity* at *p* < 0.05. D. Occipital cortex, *3O* trials: Spearman correlation (n = 16) between pN and *location recall specificity* at TRs 9-10. E. Occipital cortex; *1O1C1L* trials: Spearman correlation (n = 16) between pN and *location recall specificity* for at TR 10 in occipital cortex. F. Parietal cortex, *3O* trials: Spearman correlation (n = 16) between pN and MVPA of the empty location for TRs 6-7. G. Pearson correlation (n = 16), for *3O* trials, of parietal delay-period MVPA (of the empty location; TRs 6-7) against occipital *location recall specificity* (TRs 9-10). For D-G, solid regression lines indicate significant correlation at *p* < 0.05.

## Discussion

Varying memory load (one vs. three items) and category homogeneity on load-of-three trials (*3O* vs. *1O1C1L*) produced several novel results that provide important insight into the mechanisms underlying VWM performance. Performance in the behavior-only experiment revealed marked individual differences in swap errors, allowing us to group subjects into effective context-binding (*low swap error*) and ineffective context-binding (*high swap error*) groups. Interestingly, recall precision was comparable for these two groups, suggesting separability of processes that these measures are believed to reflect: context binding and stimulus representation, respectively. For both swap-error groups, patterns of delay-period BOLD activity in parietal and in frontal cortex were sensitive to category homogeneity rather than to memory load per se, and this trial-related variation predicted trial-related variation in the precision of behavioral recall, implicating a role for these regions in the control of inter-item interference during VWM storage. IEM of the representation of orientation was most robust in occipital cortex. Although delay-period reconstructions did not show a significant influence of inter-individual variability in context binding, recall-related reconstructions revealed markedly weaker representation of the probed item and stronger representation of non-probed items in the high-swap error group. This represents compelling neural evidence for the fact that variability in memory-retrieval processes are an important determinant of VWM performance (Unsworth et al., 2014). MVPA of the representation of stimulus location tracked the dynamic representation of location context across the trial. Stimulus location was most strongly represented in occipital cortex at encoding, and the strength of delay-period representation of location in parietal cortex predicted both neural and behavioral outcomes at recall: *location recall specificity* in occipital cortex, and the probability of making a swap error.

### Dynamics of BOLD signal intensity

The finding that delay-period activity in parietal and frontal cortex was equivalent for *1O* and *1O1C1L*, and elevated for *3O*, replicates the analogous finding in a previous study for which both the critical stimulus category (motion) and the dimension along which item-unique context varied (ordinal position) were different (Gosseries, Yu, et al., 2018). Although these results may reflect, in part, a domain-general role for parietal cortex in context binding, the present results indicate that they are also likely to reflect a role in the control of inter-item interference. This is because the pattern of “*1O* = *1O1C1L* < *3O*” was comparable for both swap-error groups. We attribute the function of this activity to the control of inter-item interference, rather than to storage processes per se (e.g., elevated inhibition among stimulus representations from the same category), because the within-subject correlations of delay-period BOLD activity with behavioral precision were only seen in parietal and frontal cortex, not in the occipital areas that play a critical role in the representation of single-feature stimuli such as oriented lines and motion (Harrison & Tong, 2009; Riggall & Postle, 2012; Serences, Ester, Vogel, & Awh, 2009). The fact that this relation between parietal delay-period activity and behavioral precision has also been seen with stimuli from a different domain (Gosseries, Yu, et al., 2018) is also consistent with a general role for parietal cortex in the control of stimulus processing in VWM.

### The influence of context binding on storage- and on recall-related processes

Although our measure of the neural representation of stimulus location is a not a direct measure of the operation of location-context binding in VWM for nonspatial stimuli, several results from our study suggest a close link between the two. One is the fact that the delay-period representation of stimulus location in parietal cortex was stronger for the *low* than the *high swap-error* group, on both *3O* and *1O1C1L* trials. A second is the fact that individual differences in delay-period parietal MVPA predicted swap errors as well as the more temporally proximal neural correlate of swap errors, *location recall specificity* in occipital cortex. Although the initial operation of binding a stimulus to its context must occur during encoding, the results summarized here illustrate that an important consequence of the strength of context binding is the efficacy with which, at the time of recall, spatial attention is focused on the location at which the probed stimulus had been presented (operationalized by *location recall specificity*). Interestingly, our results also suggest a second, partly distinct route whereby context binding influences VWM recall: *orientation recall specificity*. That is, for both *3O* and *1O1C1L* trials, recall for *high swap-error* subjects was associated with weaker representation of the orientation of the probed item and stronger representation of the orientation of unprobed items. It remains to be determined if the lack of correlation between these effects reflects truly independent factors that can be separately influenced by context binding (i.e., spatial selective attention versus item retrieval), or if this is simply a result of the fact that our measures of the neural correlates of context binding are indirect.

### Conclusion

VWM is a cognitive ability that draws on many factors, including state-like factors that vary across time (Adam, Mance, Fukuda, & Vogel, 2015), and trait-like factors that also relate to psychometric constructs, such as general fluid intelligence (Unsworth, Fukuda, Awh, & Vogel, 2014). Consistent with recent theoretical accounts(Oberauer & Lin, 2017; Schneegens & Bays, 2017) the results presented here suggest an important role for context binding in determining VWM performance. A better understanding of context binding will contribute to a better understanding of the neural processes that support, and constrain, VWM functions.

## Acknowledgement

This work was supported by National Science Foundation of China (31730038), 973 Program (2014CB846102), National Institutes of Health grant R01MH064498 (to B.R.P.).

